# Cryo-EM of α-tubulin isotype containing microtubules revealed a contracted structure of α4A/β2A microtubules

**DOI:** 10.1101/2023.03.06.531442

**Authors:** Lei Diao, Wei Zheng, Qiaoyu Zhao, Mingyi Liu, Zhenglin Fu, Xu Zhang, Lan Bao, Yao Cong

**Author notes:** Co-corresponding authors Address corresponding author: Lan Bao, Yao Cong, (L.B.); (Y.C.). These authors contributed equally to this work (Lei Diao, Wei Zheng, Qiaoyu Zhao).

## Abstract

Microtubules are hollow α/β-tubulin heterodimeric polymers playing critical roles in cells. In vertebrates, both α- and β-tubulin have multiple isotypes encoded by different genes, which are intrinsic factors in regulating microtubule functions. However, structures of microtubules composed of different tubulin isotypes, especially α-tubulin isotypes, remain largely unknown. Here, we purified recombinant tubulin heterodimers composed of different mouse α-tubulin isotypes, including α1A, α1C and α4A, with β-tubulin isotype β2A. We further assembled and determined the cryo-electron microscopy (cryo-EM) structures of α1A/β2A, α1C/β2A, and α4A/β2A microtubules. Our structural analysis demonstrated that α4A/β2A microtubules exhibit a longitudinal contraction between tubulin interdimers compared with α1A/β2A and α1C/β2A microtubules. Collectively, our findings reveal that α-tubulin isotype composition could tune microtubule structures, and also provide evidence for the “tubulin code” hypothesis.

## Introduction

Microtubules are cytoskeletal filaments playing various important roles in cellular processes, including intracellular transport, cell division, and establishment of cell polarity^1,2^. They are highly dynamic tubular polymers that assemble from α/β-tubulin heterodimers, which are arranged in a head-to-tail manner to form protofilaments, with ∼13 protofilaments associating laterally to form a hollow polar tube^3^. Both α- and β-tubulin comprise multiple genes in vertebrates, for example, mice have more than seven α- and eight β-tubulin genes, and each gene encodes a specific tubulin isotype^4^. Distinct α- and β-tubulin isotypes are expressed in different cell types to achieve diversity in microtubule organization and function^5^. The α-tubulin isotype α1A is highly expressed in post-mitotic neurons, but is decreased in postnatal and adult stages^6,7^. The β-tubulin isotype β3 is specifically expressed in neurons and important for neurite formation^8,9^. Moreover, various mutations in human tubulin genes have been reported to be closely related to diseases, especially neurodevelopmental and neurodegenerative disorders^10^, indicating that some tubulin isotypes display unique functions in these processes.

Previous studies have shown that some tubulin isotypes or tubulin constitutions could affect microtubule functions and structures. In mice, β-tubulin isotypes such as β1, β2B, and β4A did not fully rescue the neural migration defects caused by the down-regulation of neuronal β3 expression^9^. *In vitro*, purified recombinant α/β3 microtubules had a higher catastrophe frequency than α/β2B microtubules^11^. Cryo-electron microscopy (cryo-EM) studies revealed that the yeast microtubule lattice was expanded compared with mammalian microtubules^12^. Another study on *C. elegans* and *B. taurus* microtubules showed that in the absence of templates, the nucleated *C. elegans* microtubules have a smaller protofilament numbers compared to *B. taurus* microtubules^13^. Further, it has been suggested that the lateral contact loops are ordered in *C. elegans* but unresolved in *B. taurus*^13^. Moreover, other works with single tubulin isotype purified from insect cells showed that GMPCPP-α1A/β3 microtubules have subtle differences at polymerization interfaces compared with GMPCPP-brain microtubules^14^, and GMPCPP-α1B/β2B microtubules are more stable and consist of more protofilaments than α1B/β3 ones^5^. Altogether, it appears that microtubules assembled with different tubulin isotypes or tubulin constitutions exhibit distinct microtubule structures. There is also a “tubulin code” hypothesis stated that multiple tubulin isotypes and diverse post-translational modifications determine microtubule properties and functions^15^.

The α/β-tubulin heterodimers are composed of α- and β-tubulin isotypes, and each subunit contains a GTP binding site at the longitudinal interface between the isotypes. The GTP bound to β-tubulin, at the E-site (exchangeable), is hydrolyzed within the microtubule assembly proceeding^16^ via longitudinal contacts with α-tubulin, while the GTP bound to α-tubulin, at the N-site (non-exchangeable), plays a structural role and is never hydrolyzed^17^. Recently, we reported that α1A/β2A and α1C/β2A form microtubules displaying distinct properties, which was mainly mediated by the C-terminal tail of α-tubulin^18^. However, the effects of specific α-tubulin isotype on microtubules structure remains largely unknown.

Here, we sort to examine the structural features of α1A/β2A and α1C/β2A microtubules, exhibiting large distinct dynamics^18^, and another α4A/β2A microtubules, possessing a special bending feature^19^. By using total internal reflection fluorescent (TIRF) microscopy assay, we showed that α1A/β2A, α1C/β2A and α4A/β2A tubulin dimers are all able to assemble into microtubules in the presence of GMPCPP and taxol. We determined the cryo-EM structures of the α1A/β2A, α1C/β2A, and α4A/β2A microtubules at the resolution range of 4.2 Å to 4.4 Å. Furthermore, compared to α1A/β2A and α1C/β2A microtubules, α4A/β2A microtubules display a longitudinal contraction between tubulin interdimer, suggesting that α-tubulin isotypes are able to tune microtubule structure. This study also expands the current knowledge of α-tubulin isotypes on microtubule functions.

## Results

### Recombinant mouse α1A/β2A, α1C/β2A, and α4A/β2A form microtubules

The amino-acid sequences of α-tubulin isotypes are highly conserved. Our recent work on α1A/β2A, α1C/β2A and α4A/β2A microtubules suggested that microtubule polymerization properties and morphologies could be affected by specific α-tubulin isotypes^18,19^. To further explore the effect of α-tubulin isotypes on microtubule structures, we expressed and purified different α-tubulin isotypes, including α1A/β2A, α1C/β2A, and α4A/β2A, using a previously established method (Figures 1A and 1B)^18^. TIRF assay showed that α1A/β2A, α1C/β2A, and α4A/β2A tubulin dimers are all able to assemble into microtubules in the presence of GMPCPP or taxol (Figures 1C and 1D). Moreover, direct visualization of the cryo-EM images showed that these three types of tubulin dimers could all form microtubules, with which decorated with a kinesin mutant K_349_(E236A) (Figures 1E and 1F). Putting together, tubulin dimers composed of an α-tubulin isotype (including α1A, α1C and α4A) and a constant β-tubulin β2A are able to form microtubules.

**Figure 1.**
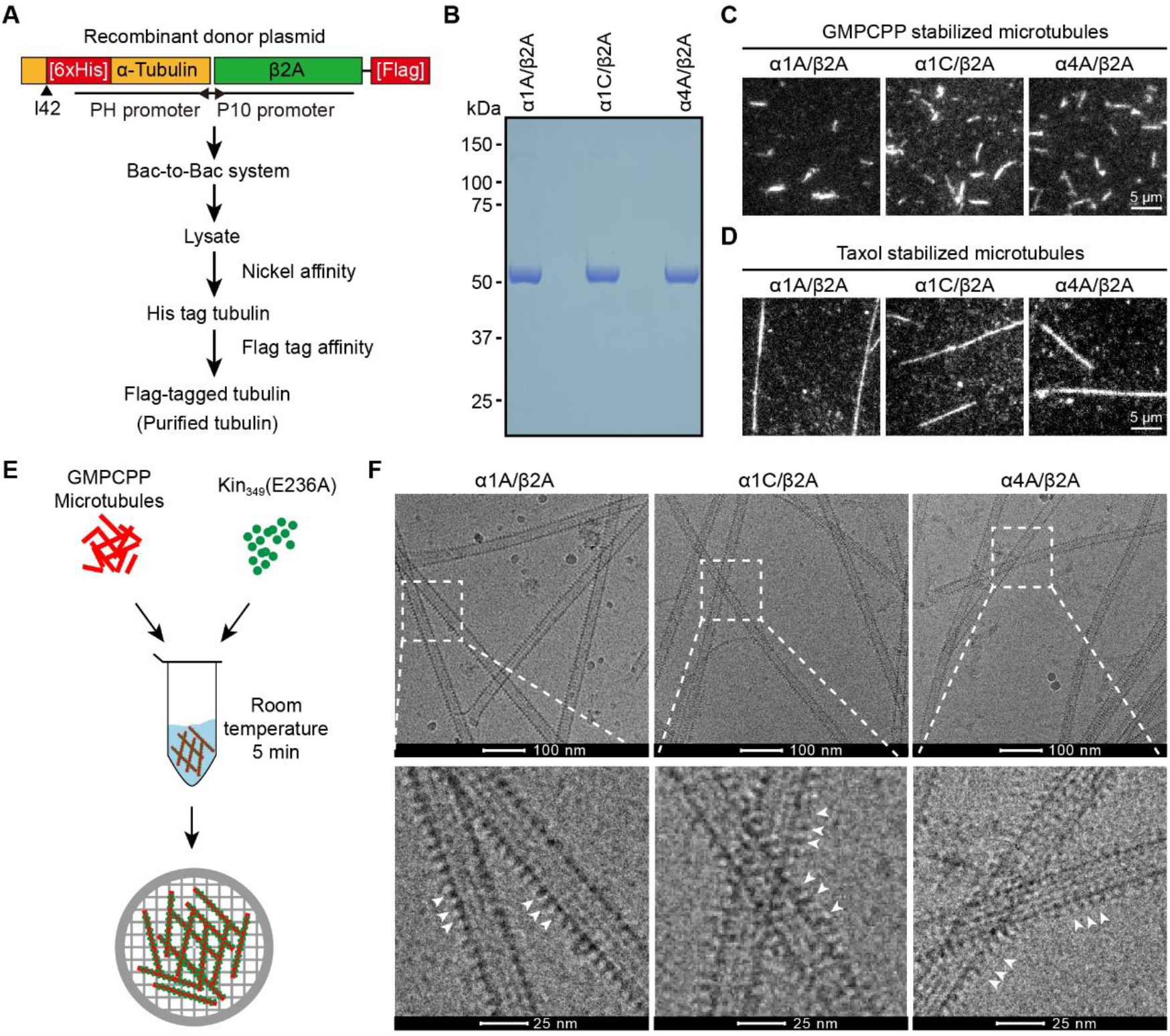
Purification of recombinant mouse α/β-tubulin dimers. (A) Schematic diagram for purification of recombinant tubulin isotypes. (B) Coomassie blue staining of purified α1A/β2A, α1C/β2A, and α4A/β2A. (C and D) TIRF images of GMPCPP (C) and Taxol (D) stabilized α1A/β2A, α1C/β2A, and α4A/β2A microtubules. Rhodamine-labeled porcine tubulin (∼4%) was added to visualize microtubules, and biotin-labeled porcine tubulin (∼4%) was added to immobilize microtubules on neutravidin-coated coverslips. (E) Schematic for cryo-EM sample preparation of α1A/β2A, α1C/β2A, and α4A/β2A microtubules decorated with Kin_349_(E236A). (F) Representative cryo-EM images of α1A/β2A, α1C/β2A, and α4A/β2A microtubules decorated with Kin_349_(E236A) (indicated by white arrow head).

### Cryo-EM structures of α1A/β2A, α1C/β2A, and α4A/β2A microtubules

To investigate the architectures of α1A/β2A, α1C/β2A, and α4A/β2A microtubules and explore the impact of α-tubulin isotypes on the structural features of the microtubules, we characterized the structures of the three kinds of microtubules in the presence of GMPCPP by using cryo-EM. The microtubules were decorated with K_349_(E236A) to aid in the identification of α- and β-tubulin in the reconstruction process through the “seam-search” protocol (Figures 1E and 1F)^3,20^. We determined the cryo-EM maps of α1A/β2A, α1C/β2A, and α4A/β2A microtubules in the dominantly populated 14-protofilament type at the resolution range of 4.2 Å to 4.4 Å (Figures 2A-2C; Figure S1; Table S1; Methods). We then built the corresponding atomic model for each of the maps, including the non-seam region and the seam region (Figure 2D; Figures S2A and S2B).

**Figure 2.**
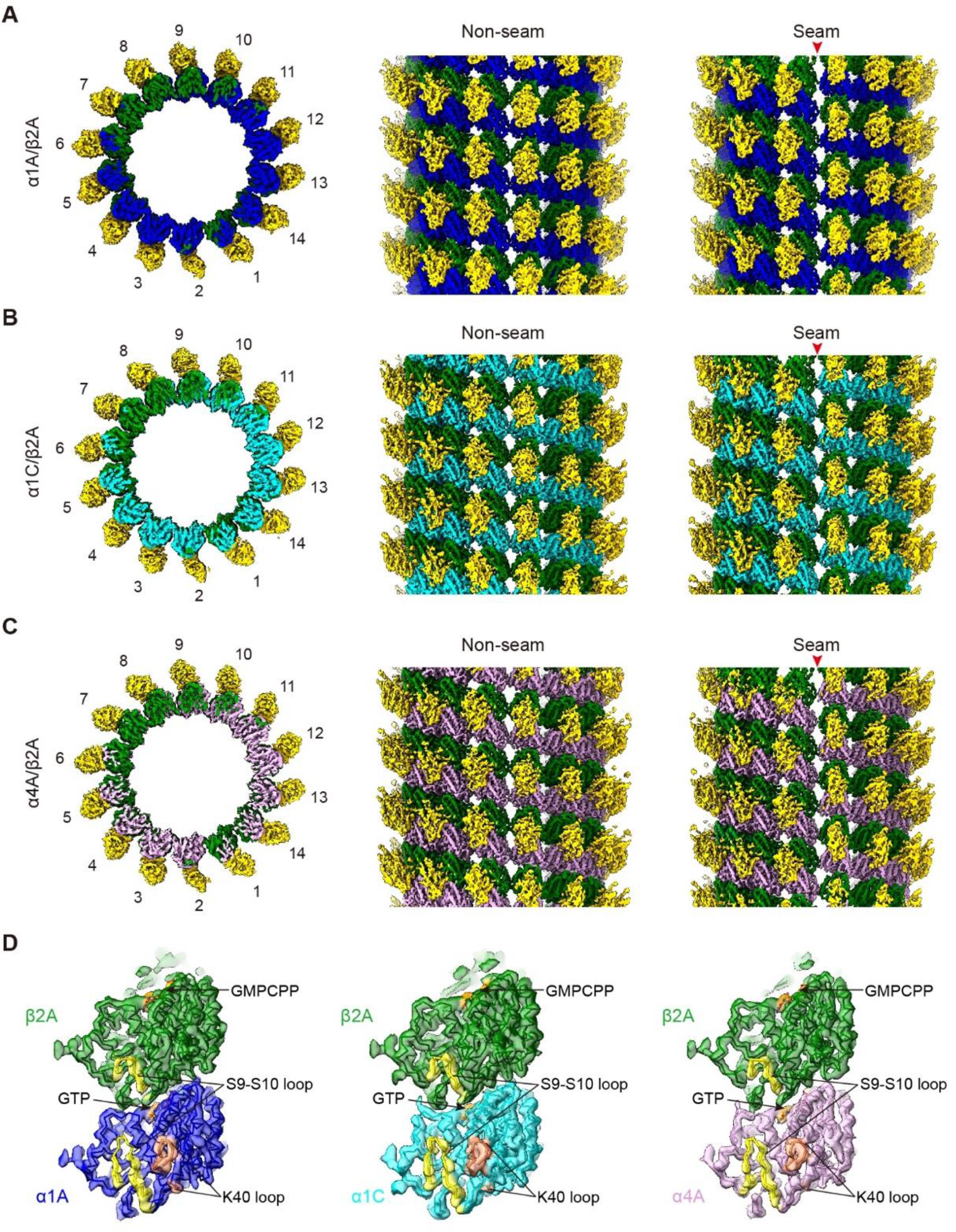
Cryo-EM structures of α1A/β2A, α1C/β2A, and α4A/β2A microtubules. (A-C) Overview of the cryo-EM map for α1A/β2A (A), α1C/β2A (B), and α4A/β2A (C) microtubules decorated with kinesin-1 motor domain, respectively. Kinesin is in gold, β2A in green, α1A in blue, α1C in cyan, and α4A is pink; this color scheme is followed throughout. Viewed from the top and side of microtubules, respectively. (D) Model and map (transparent surface) fitting of α1A/β2A, α1C/β2A, and α4A/β2A microtubules. Viewed from the lumen side, with the key structural elements indicated.

Both a- and β-tubulin isotypes contain a S9-S10 loop, which is eight residues longer in α-tubulin than that in β-tubulin and forms part of the taxol-binding site in β-tubulin^21,22^. Here in our α1A/β2A, α1C/β2A, and α4A/β2A structures, it appears that the S9-S10 loop in α-tubulin is indeed longer than that in β-tubulin (Figure 2D), demonstrating a successful separation of α- and β-tubulin in these reconstructions. Still, the density of α-tubulin acetylation loop P37-G45 (also known as K40 loop) and the C-terminal tail of both α- and β-tubulin were not observed in all maps (Figure 2D), likely due to the well-known flexibility of these regions^22^.

### Similar lateral contacts among α1A/β2A, α1C/β2A, and α4A/β2A microtubules

To understand the effects of α-tubulin on microtubule structure, we first analyzed the structures of α1A/β2A, α1C/β2A, and α4A/β2A dimers and found that these dimers exhibit overall similar conformation (Figures S2C and S2D). We then compared the models of two tubulin dimers within adjacent protofilaments in the three obtained microtubule structures. By superimposing the global Cα of the α-tubulin on the left side, we found that for α1C/β2A microtubules, the lateral interaction between α-α and β-β contact in the non-seam region was similar to that of α1A/β2A ones, with the Cα atom root mean square deviation (RMSD) lower than 1 Å (Figure 3A). This was also the case for α4A/β2A and α1A/β2A microtubules (Figure 3B). Similar analysis on the lateral interaction between α-β and β-α contact in the seam region suggested that the α1C/β2A microtubules display a subtle longitudinally contracted divergence between the α1C and α1A in the seam region (Figure 3C); the α4A/β2A microtubules also display a slight longitudinal contraction (Cα atom RMSD of ∼1 Å) between the α4A and α1A in the seam region (Figure 3D). Also, we did not observe obvious structural deviation in the lateral interfaces of these three types of microtubules (Figures 3A-3D). Taken together, our data suggest that these α-tubulin isotypes don’t show obvious lateral effects on their microtubule structures, while α1C/β2A and α4A/β2A microtubules exhibit slight longitudinal contraction between the interdimers in the seam region.

**Figure 3.**
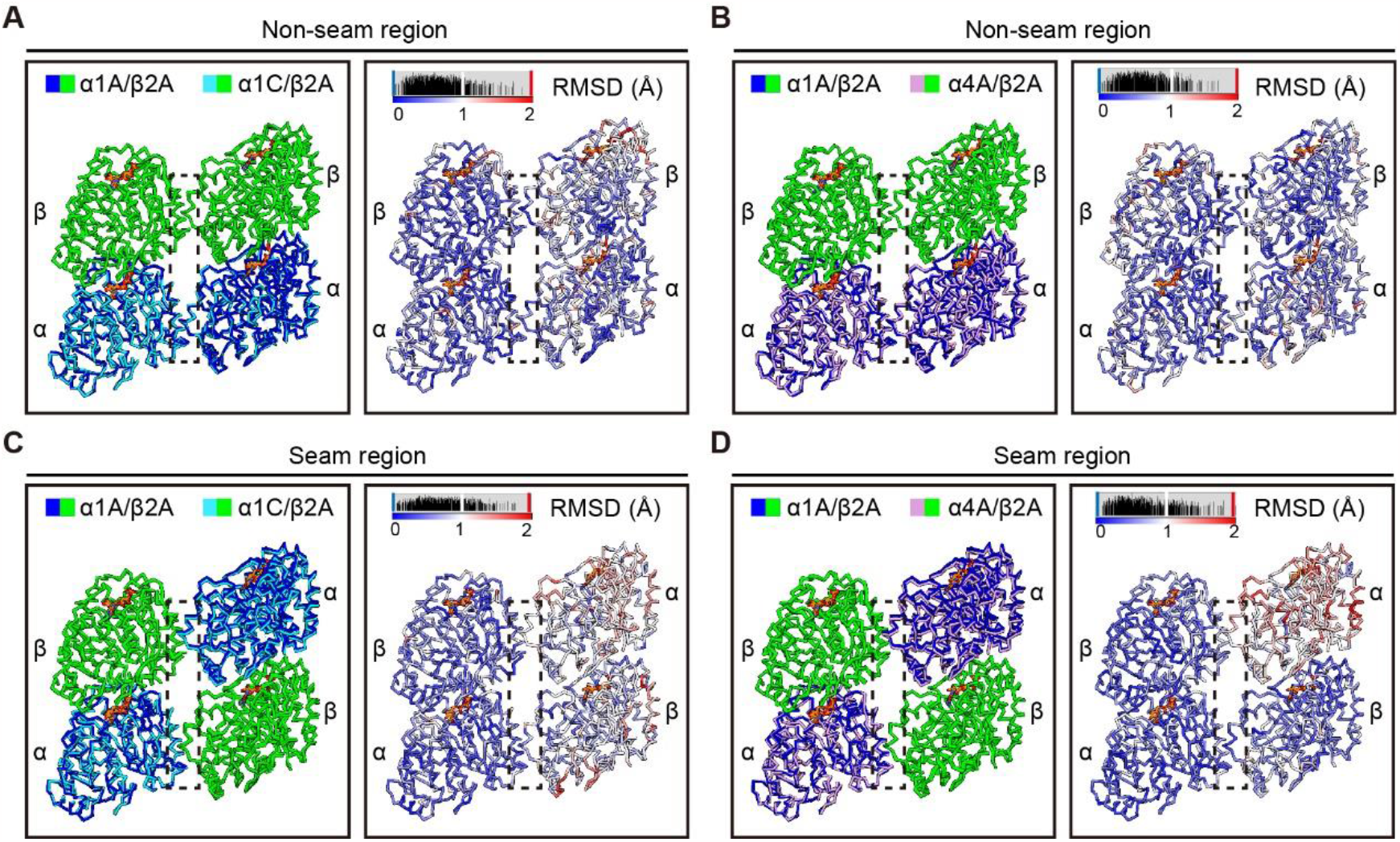
The lateral interactions are similar among α1A/β2A, α1C/β2A, and α4A/β2A microtubules. (A, B) Visualization of the lateral interactions in the non-seam region, by comparison of the atomic models between α1A/β2A and α1C/β2A microtubules (A) and between α1A/β2A and α4A/β2A microtubules (B), and corresponding Cα atom RMSDs. (C, D) Visualization of the lateral interactions in the seam region, by comparison of the atomic models between α1A/β2A and α1C/β2A microtubules (C) and between α1A/β2A and α4A/β2A microtubules (D), and corresponding Cα atom RMSDs. Viewed from the lumen side. The lateral interaction interface between the neighboring protofilament is indicated by dotted black frame.

### A distinct contracted lattice of α4A/β2A microtubules

Due to the longitudinally contracted divergence in the seam region, especially between α1A/β2A and α4A/β2A microtubules (Figure 3D), we further compared the structures of α1A/β2A and α1C/β2A, and that of α1A/β2A and α4A/β2A in the longitudinal orientation. We first superimposed three consecutive tubulin dimers within a single protofilament of α1C/β2A and α1A/β2A microtubules by aligning the Cα of the bottom α-tubulin together. We found that the longitudinal interactions between intradimer and interdimer of α1A/β2A and α1C/β2A microtubules are very similar (Figures 4A-4E), which was confirmed by Cα atom RMSD analysis (Figure 4B). Surprisingly, similar analysis on α1A/β2A and α4A/β2A microtubules revealed a clear contraction of α4A/β2A relative to the α1A/β2A protofilament (Figures 4F-4J). This contraction mainly exists between the longitudinal interdimer, consistent with that observed in the seam region (Figures 3C and 3D). Also, this contraction is in a progressive expanded manner that increased with each dimer along the protofilament, such that the first neighboring α4A/β2A dimer contracted by ∼1.5 Å relative to α1A/β2A microtubules, then the second dimer by 3 Å, and so on. Cα atom RMSD analysis also confirmed these contraction divergences (Figure 4G). Collectively, our structural analysis revealed that α4A/β2A microtubules displayed a longitudinal contraction between tubulin interdimer compared with α1A/β2A and α1C/β2A microtubules, indicating that microtubules structures can be regulated by different α-tubulin isotypes.

**Figure 4.**
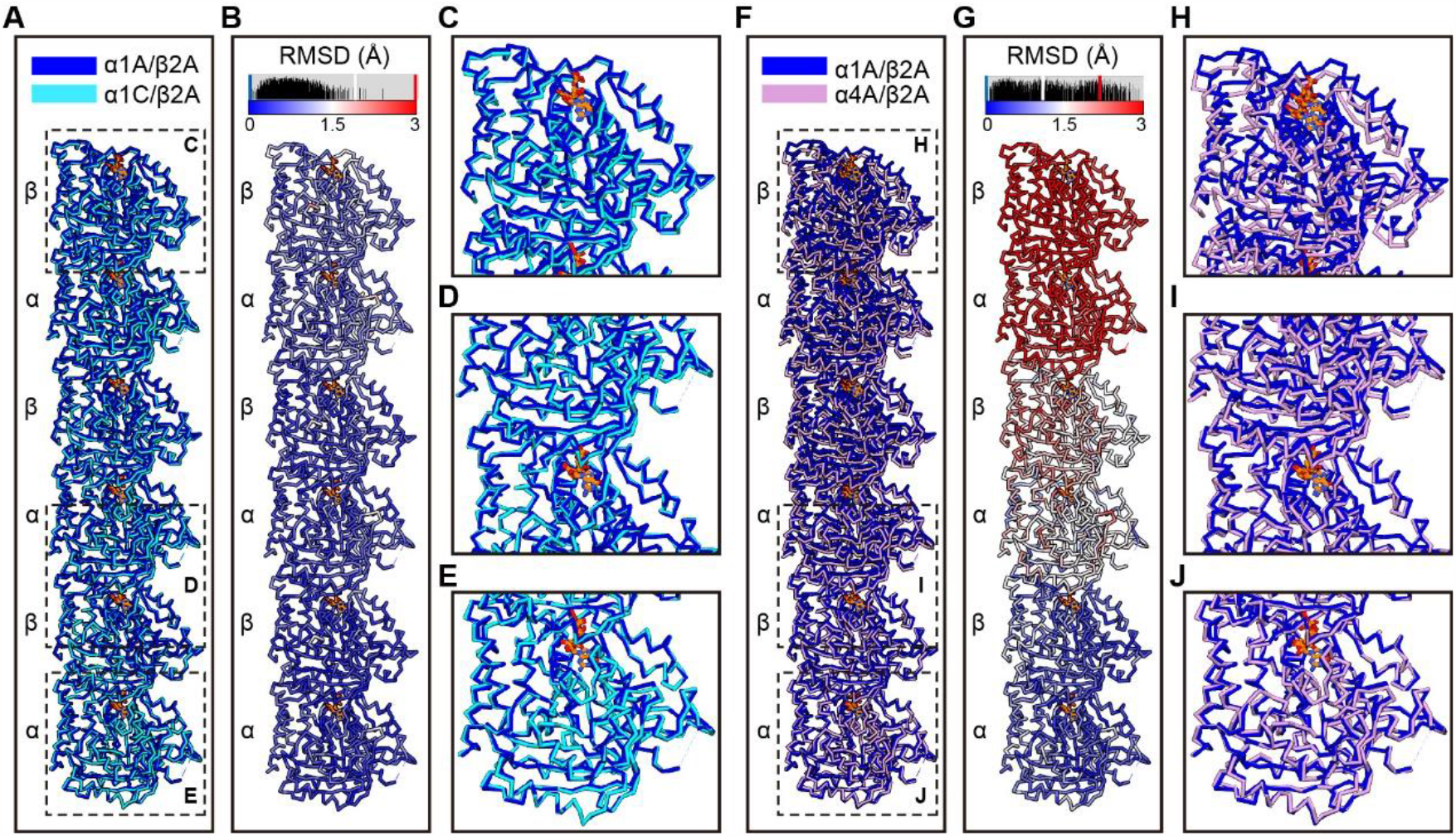
α4A/β2A microtubules display a longitudinal contracted lattice. (A) Comparison of the Cα trace of three consecutive tubulin dimers between α1A/β2A and α1C/β2A microtubules, superimposed on the bottom α-tubulin. (B) Cα atom RMSDs between the two models shown in (A), with deviations colored from blue to red. (C-E) Zoom-in view of the boxed regions in (A). Viewed from the lumen side. The rendering style was followed throughout this figure. (F) Comparison of the Cα trace of three consecutive tubulin dimers between α1A/β2A and α4A/β2A microtubules, superimposed on the bottom α-tubulin. (G) Cα atom RMSDs between the two models shown in (F). (H-J) Zoom-in view of the boxed regions in (F).

## Discussion

Microtubules assembled from different tubulin constitutions or with different tubulin isotypes exhibit distinct microtubule structures. For instance, previous cryo-EM studies revealed that microtubules assembled from different sources, such as yeast, porcine brain, *C. elegans* or *B. taurus* brain, exhibit different structures^12,13,23^. There were also other cryo-EM studies with single tubulin isotypes showed that GMPCPP-α1A/β3 and GMPCPP-brain microtubules display different microtubule structures^14^, which was also the case for GMPCPP-α1B/β2B and GMPCPP-α1B/β3 microtubules^5^. In the current study, we assembled and determined the cryo-EM structures of α1A/β2A, α1C/β2A, and α4A/β2A microtubules, not available before. Our structural analysis revealed that, compared with α1A/β2A and α1C/β2A microtubules, α4A/β2A microtubules exhibit a longitudinal contraction between tubulin interdimers, which could potentially be related to the more amino acids variations between α4A and α1A relative to that between α1C and α1A (Figures S2C and S2D). Collectively, our results suggest that α-tubulin isotypes can tune microtubules structure.

Eukaryotic cells usually contain multiple α- and β-tubulin isotypes and their expression patterns vary widely among different tissues and during different developmental stages^24,25^. Tubulin isotype α4A is highly expressed in various tissues such as heart, brain and platelet^26,27^, and is mainly expressed at later stages of development^28^. Previous studies have suggested that some mutations on tubulin α4A could cause serious diseases, including amyotrophic lateral sclerosis (ALS)^29-31^, frontotemporal dementia (FTD)^32^, and macrothrombocytopenia^27^. Moreover, it has been reported that tubulin isotype β2 is predominantly present in the neurites of differentiated neuroblastoma SK-N-SH cells and plays a key role in neurite outgrowth, while β1 and β3 both occur in cell bodies and neurites^33^. However, the distribution of most tubulin isotypes in cells remains largely unclear, mainly due to the lack of specific antibodies to distinguish these highly conserved isotypes. Here we found that α4A/β2A microtubules display a contracted structure, which provides powerful evidence that different tubulin isotype constitutions could alter the structural properties of microtubules. Collectively, our previous^18,19^ and current studies demonstrate that α-tubulin isotype composition can tune microtubules dynamics, morphology and structure. Still, the functional role of specific tubulin isotype, such as α4A, in cells remains to be further explored.

## Materials and Methods

### Plasmid construction

The expression constructs for α1A (NM_011653.2), α1C (NM_009448.4), α4A (NM_009447.4), and β2A (NM_009450.2) were cloned from cDNA of mouse brain tissue into the vector pFastBac Dual. α1A, α1C, and α4A sequences were inserted after the polyhedron promoter, and β2A sequence was inserted after the p10 promoter. For affinity purification, a sequence encoding GGSGG linker and a Flag tag were fused to the 3’ end of β2A sequence. For α1A, α1C, and α4A, a His tag was inserted in the acetylation loop (between I42 and G43)^34^. An enhancer L21^35^ was added to each of these sequences just before the start codon. The expression construct for kinesin motor domain (KIF5B, NM_004521.3), truncated (amino acids 1-349) and defective in ATP hydrolysis (E236A) ability [Kin_349_(E236A)], was cloned from cDNA of HEK293T cells into vector pET28a with a His tag before the N-terminal of Kin_349_(E236A).

### Expression and purification of mouse recombinant tubulin

Recombinant tubulin was purified by using Bac-to-Bac system (Life Technologies) as described previously^18^. Briefly, SF9 cells (Life Technologies) were grown to 2.0-2.5× 10^6^ cells/ml in Sf-900™ II SFM (Thermo Fisher 10902088) supplemented with antibiotics including penicillin and streptomycin and infected with P3 viral stocks. Cells were cultured in suspension at 27°C and harvested at 72 h after infection. The following steps were performed at 4°C or on ice. Cells were lysed by sonication in lysis buffer (80 mM PIPES pH 6.9, 100 mM KCl, 1 mM MgCl_2_, 1 mM EGTA, 0.1 mM GTP, 0.5 mM ATP, 1 mM PMSF), and then the lysate was centrifuged for 30 min at 35,000 g. The supernatant was filtered through a 0.45 μm Millex-HV PVDF membrane and loaded on a nickel-nitrilotriacetic acid column (Qiagen) pre-equilibrated with lysis buffer. The column was washed with 30 ml wash buffer (lysis buffer supplemented with 25 mM imidazole), and then eluted with elution buffer [1× BRB80 (80 mM PIPES, 1 mM MgCl_2_, 1 mM EGTA), 300 mM imidazole, 0.1 mM GTP, pH 7.0]. The eluate was diluted with an equal volume of BRB80 buffer supplemented with 0.1 mM GTP, and mixed with anti-Flag antibody-conjugated resin (Sigma-Aldrich) for 2 h. Flag-tagged tubulin was eluted with BRB80 buffer supplemented with 0.2 mg/ml 3× Flag peptide (APE×BIO A6001). Finally, the purified tubulin was concentrated and desalted on an Amicon Ultracel-30 K filter (Milipore, Merck KGaA) with BRB80 and 0.1 mM GTP. The tubulin concentration was estimated by UV at 280 nm, frozen in liquid nitrogen and stored at -80°C.

### Purification of Kin_349_(E236A)

The Kin_349_(E236A) was expressed and purified from *E. coli* BL21 (DE3) using a protocol that was modified from the previously published methods^36^. The transformed *E. coli* BL21 (DE3) cells were cultured in Luria-Bertani medium (LB) at 37°C until the OD_600nm_ was between 0.6 and 0.8, and then were added 0.5 mM isopropyl-β-D-thiogalactopyranoside (IPTG, Sigma) for the induction of protein expression at 20°C for 16-18 h. Cells were harvested and resuspended in lysis buffer (20 mM PIPES pH 6.9, 150 mM KCl, 4 mM MgCl_2_, 0.1 mM ATP, 1 mM PMSF) and lysed by sonication. The lysate was cleared by centrifugation at 35,000 g for 20 min at 4°C. The supernatant was filtered through a 0.45 μm filter and loaded on a nickel-nitrilotriacetic acid column (Qiagen) pre-equilibrated with lysis buffer. The column was washed with 30 ml wash buffer and eluted with elution buffer (80 mM PIPES pH 6.9, 4 mM MgCl_2_, 1 mM EGTA, 300 mM imidazole, 0.1 mM ATP). The proteins were concentrated and gel-filtrated on a SuperdexTM 75 size exclusion chromatography column (GE Healthcare) equilibrated with stock buffer (80 mM PIPES pH 6.9, 100 mM KCl, 4 mM MgCl_2_, 1 mM EGTA, 0.1 mM ATP). Protein fractions containing the target protein were collected and concentrated. The protein concentration was estimated by UV at 280 nm, frozen in liquid nitrogen and stored at -80°C.

### Microtubule preparation for TIRF assay

For the GMPCPP or taxol stabilized α1A/β2A, α1C/β2A, and α4A/β2A microtubules, 7.5 μM of α1A/β2A, α1C/β2A, or α4A/β2A tubulins ∼4% biotin-tubulin (Cytoskeleton) and ∼4% rhodamine-tubulin (Cytoskeleton) were incubated with 1 mM GMPCPP or 1 mM GTP with 10 mM taxol at 37°C for 3 h, respectively. Then, the solution was diluted in warmed BRB80 (for GMPCPP microtubules) or BRB80 containing 1 mM GTP and 10 mM taxol (for taxol microtubules) and incubated in the cleaning and silanizing glass coverslips for 5 min. Images were acquired by TIRF microscopy with Zeiss cell observer spinning disk system and a 100× oil lens. The images were recorded with a pixel size of 160 nm.

### Formation of kinesin-decorated α1A/β2A, α1C/β2A, and α4A/β2A microtubules

For the construction of the GMPCPP stabilized α1A/β2A, α1C/β2A, and α4A/β2A microtubules, 15∼20 μM of α1A/β2A, α1C/β2A, or α4A/β2A tubulins was incubated with 1 mM GMPCPP at 37°C for 3 h, respectively. To remove unpolymerized tubulin, the solution was centrifuged at 126,000 g for 5 min at 27°C and the supernatant was discarded. Then, the microtubule pellet was resuspended in cold BRB80 and depolymerized at 4°C for 20 min, followed by a second round of polymerization at 37°C with 1 mM GMPCPP for 3 h. GMPCPP-stabilized microtubules were pelleted as above, resuspended in 37°C pre-warmed BRB80 with ∼80% volume of initial reaction. The microtubule solution was diluted at a ratio of 1:2∼1:3, followed by mixing with ∼20 μM Kin_349_(E236A) and 2 mM ATP, and then incubated for 5 min at room temperature.

### Cryo-EM sample preparation

To prepare the cryo-EM sample of the kinesin-decorated α1A/β2A microtubules, an aliquot of 2.2 μl sample was applied to the plasma-cleaned holey carbon grid (Quantifoil R1.2/1.3, Cu, 400 mesh). The grid was blotted using a Vitrobot Mark IV (Thermo Fisher Scientific), using a blot force of -1 and 1 s blot time at 100% humidity and 22°C, and then plunged into liquid ethane cooled by liquid nitrogen. For the cryo-EM sample preparation of the kinesin-decorated α1C/β2A or α4A/β2A microtubules, similar procedure was applied.

### Cryo-EM data collection

Cryo-EM movies of the samples were collected on a Titan Krios transmission electron microscope (Thermo Fisher Scientific) operated at an accelerating voltage of 300 kV. All movies were recorded on a K2 Summit direct electron detector (Gatan) operated in the super resolution mode at a nominal magnification of 18,000× (yielding a pixel size of 1.3 Å after 2 times binning) under a low-dose condition^37^. Each movie was dose-fractioned into 38 frames with a dose rate of 8 e^−^ per pixel per sec on the detector. The exposure time was 7.6 s with 0.2 s for each frame, generating a total dose of 36 e^−^/Å^2^. Defocus values varied from -0.8 to -1.5 μm. All of the images were collected by utilizing the SerialEM automated data collection software package^37^.

### Image processing and 3D reconstruction

For each dataset, the motion correction of the image stack was performed using MotionCor2^38^, and CTF parameters were determined using CTFFIND4^39^. The subsequent image processing for microtubule reconstruction was performed mainly following the previously established method and related procedure^3^. Briefly, EMAN1 program *helixboxer* was used to select microtubules^40^. Subsequently, the microtubules were divided into overlapping microtubule “particles”, with the box size of 512 pixels and the non-overlapping distance of adjacent microtubule particles set to ∼80 Å. Multi-reference alignment (MRA) in EMAN1 was used to determine the initial alignment parameters and protofilament number^40^. The microtubule particles with the same protofilament number were merged and subjected to further refinement in FREALIGN v9^41^. The number of microtubule protofilaments formed *in vitro* varies from 9 to 16, with the majority consisting of 14 protofilaments^42^. Indeed, over 55% of the microtubules contained 14 protofilaments in our assay. Thus, only the dominant 14-protofilament microtubule densities were used for further refinement, with a seam-search protocol described previously^20^. The final resolution for each reconstruction was estimated based on the gold-standard criterion using a Fourier shell correlation (FSC) of 0.143. All of the reconstructions were post-processed through deepEMhancer^43^, and the reconstructions were sharpened by applying a b-factor value of -100 Å^2^. UCSF Chimera and ChimeraX were applied for structural visualization and figure generation^44^.

### Atomic model building and analysis

Initial models of α1A/β2A, α1C/β2A, and α4A/β2A microtubules were generated through the SWISS-MODEL server^45^, using an existing GMPCPP-bound *Chlamydomonas reinhardtii* tubulin model (PDB 6U42) as the template. For the seam region, 16 adjacent tubulin dimer models were fit into the corresponding region of the map in Chimera by rigid-body fitting, which were then refined against the corresponding map by appling *phenix*.*real_space_refine* in Phenix^46^, to make sure the model of the central 4 dimers to match well with the density map. For the non-seam region, 9 dimer models were used to perform the model refinement following similar procedure, to ensure that the model of the central dimer to match well with the density map. All RMSD calculations were performed using UCSF Chimera^44^.

## Acknowledgements

We thank Dr. Rui Zhang at Washington University in St. Louis, for his instruction in microtubule structure determination. We are grateful to the staffs of the Electron Microscopy facility, Laser Microscopy facility, Database and Computing facility, and Mass Spectrometry facility at National Facility for Protein Science in Shanghai (NFPS), Zhangjiang Lab, China for instrument support and technical assistance. Our work was supported by grants from the National Natural Science Foundation of China (31991194, 31330046, and 32130056), the Strategic Priority Research Program of Chinese Academy of Sciences (XDB19000000, XDB37040103), National Key R&D Program of China (2017YFA0503503), Shanghai Academic Research Leader (20XD1404200), and Shanghai Pilot Program for Basic Research from CAS (JCYJ-SHFY-2022-008).

## Author contributions

L.B. and Y.C. supervised the entire project. L.D. purified the proteins and performed biochemical studies. M.L. involved in protein purifications. X.Z. helped with the experiment design. W.Z. performed the cryo-EM data acquisition with the involvement of Z.F. W.Z. performed 3D reconstruction and structural analysis with the assistance of Q.Z. L.D., Y.C., and L.B. wrote the manuscript with the input of W.Z.

## Data availability

All data presented in this study are available within the figures and the supplementary information. Cryo-EM maps have been deposited in the EMDB (ID code ***, *** and ***), and the associated models have been deposited in the PDB (ID code ***, *** and ***).

## Supplementary

**Figure S1.**
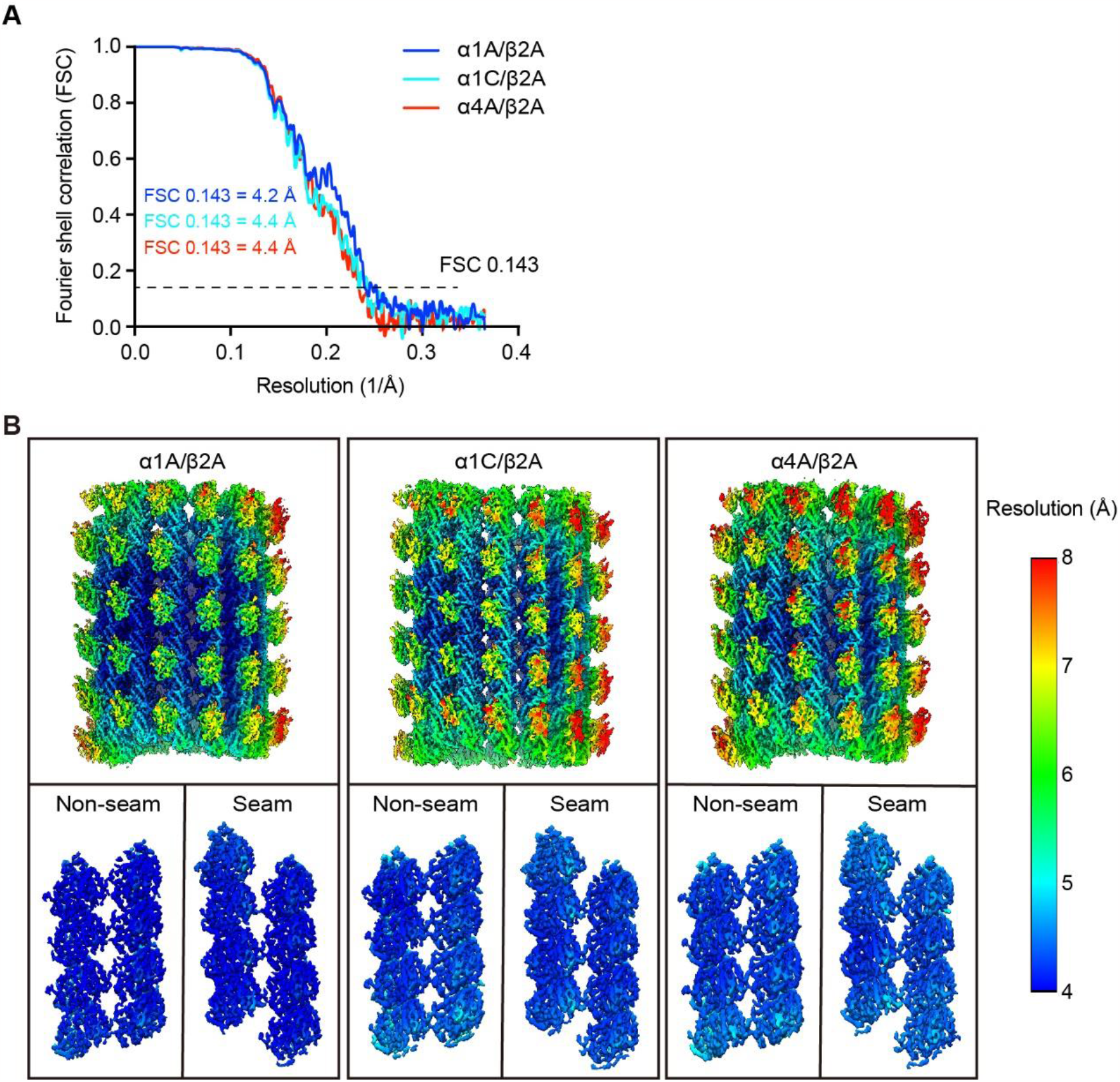
Cryo-EM reconstructions of GMPCPP-stabilized α1A/β2A, α1C/β2A, and α4A/β2A microtubules decorated with Kin_349_(E236A). (A) Resolution estimation of the cryo-EM maps for α1A/β2A, α1C/β2A, and α4A/β2A microtubules according to the gold-standard FSC criterion of 0.143. (B) Local resolution estimation of the α1A/β2A, α1C/β2A, and α4A/β2A microtubule maps, with their non-seam and seam regions also displayed from the lumen side. The resolution color bar is shown on the right.

**Figure S2.**
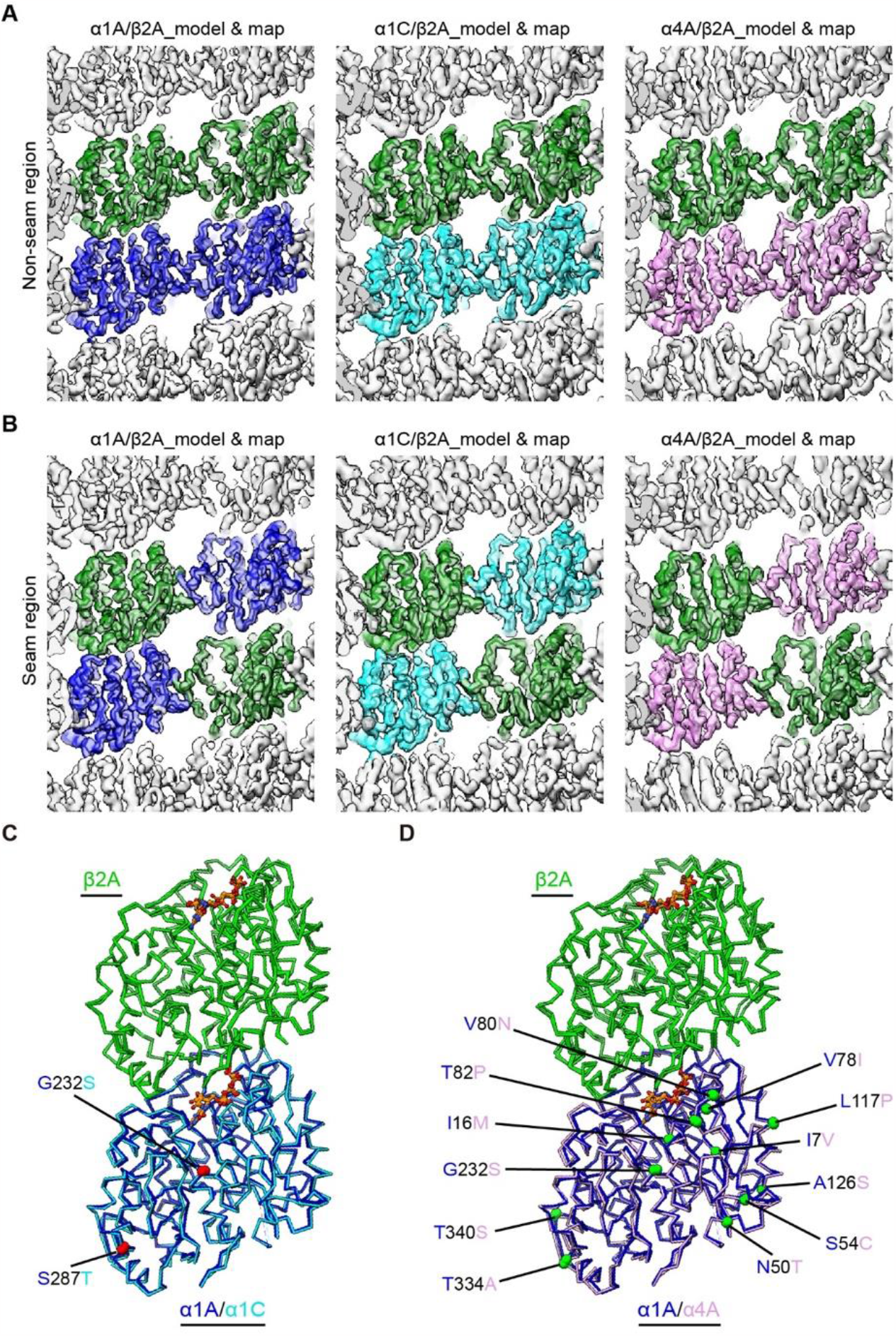
Cryo-EM density maps and models of α1A/β2A, α1C/β2A, and α4A/β2A microtubules. (A and B) Model and map fitting of α1A/β2A, α1C/β2A, and α4A/β2A microtubules, showing homotypic lateral interactions at the non-seam region (A) and at the seam region (B), viewed from the lumen side. (C) Comparison of the Cα trace for tubulin dimers between α1A/β2A and α1C/β2A microtubules. The distinct amino acids between α1A and α1C were highlighted in red balls. (D) Comparison of the Cα trace for tubulin dimers between α1A/β2A and α4A/β2A microtubules. The distinct amino acids between α1A and α4A were highlighted in green balls. It appears there are more amino acids variations between α4A and α1A (D) relative to that between α1C and α1A (C).

**Table S1.** Cryo-EM data collection and refinement statistics.

| Sample | $\alpha 1A/\beta 2A$ | $\alpha 1C/\beta 2A$ | $\alpha 4A/\beta 2A$ |
| --- | --- | --- | --- |
| **Data collection** |  |  |  |
| EM equipment | Titan Krios | Titan Krios | Titan Krios |
| Voltage (kV) | 300 | 300 | 300 |
| Detector | K2 Summit | K2 Summit | K2 Summit |
| Pixel size (Å) | 1.3 | 1.3 | 1.3 |
| Total electron dose (e<sup>-</sup>/Å<sup>2</sup>) | 36 | 36 | 36 |
| Dose rate (e<sup>-</sup>/physical pixel/s) | 8 | 8 | 8 |
| Exposure time (s) | 7.6 | 7.6 | 7.6 |
| Defocus range (μm) | -0.8 ~ -1.5 | -0.8 ~ -1.5 | -0.8 ~ -1.5 |
| **Reconstruction** |  |  |  |
| Software | EMAN & Frealign | EMAN & Frealign | EMAN & Frealign |
| Original particles | 32,977 | 31,952 | 33,817 |
| Final particles (14 protofilaments) | 19,287 | 21,250 | 24,668 |
| Final particles (13 protofilaments) | 7,068 | 5,228 | 4,845 |
| Symmetry | Pseudo-Helix | Pseudo-Helix | Pseudo-Helix |
| Final resolution (Å) | 4.2 | 4.4 | 4.4 |
| Twist (°) | -25.75 | -25.75 | -25.78 |
| Rise (Å) | 8.72 | 8.72 | 8.62 |
| Map sharpening B-factor | -100 | -100 | -100 |
| **Atomic modeling (Seam)** |  |  |  |
| Software | SWISS-MODEL & Phenix | SWISS-MODEL & Phenix | SWISS-MODEL & Phenix |
| Model composition |  |  |  |
| Atoms | 148,380 | 148,396 | 148,364 |
| Protein residues | 18,848 | 18,848 | 18,848 |
| Ligands | 48 | 48 | 48 |
| Rms deviation |  |  |  |
| Bond length (Å) | 0.006 | 0.005 | 0.005 |
| Bond angle (°) | 1.17 | 0.91 | 1.17 |
| Ramachandran plot |  |  |  |
| Favored (%) | 97.32 | 97.59 | 96.91 |
| Allowed (%) | 2.68 | 2.41 | 3.09 |
| Outliers (%) | 0.00 | 0.00 | 0.00 |
| Clash score | 2.86 | 4.01 | 3.57 |
| **Atomic modeling (Non-seam)** |  |  |  |
| Model composition |  |  |  |
| Atoms | 84,105 | 84,114 | 84,105 |
| Protein residues | 10,602 | 10,602 | 10,602 |
| Ligands | 27 | 27 | 27 |
| Rms deviation |  |  |  |
| Bond length (Å) | 0.005 | 0.005 | 0.005 |
| Bond angle (°) | 1.17 | 1.17 | 1.18 |
| Ramachandran plot |  |  |  |
| Favored (%) | 97.44 | 97.76 | 97.92 |
| Allowed (%) | 2.56 | 2.24 | 2.08 |
| Outliers (%) | 0.00 | 0.00 | 0.00 |
| Clash score | 3.85 | 3.33 | 3.35 |

